# Sensitivity to gains during risky decision-making differentiates chronic cocaine users from stimulant-naïve controls

**DOI:** 10.1101/795443

**Authors:** B. Kluwe-Schiavon, A. Kexel, G. Manenti, D.M. Cole, M.R. Baumgartner, R. Grassi-Oliveira, P.N. Tobler, B.B. Quednow

## Abstract

**Background:** Although chronic cocaine use has been frequently associated with decision-making impairments that are supposed to contribute to the development and maintenance of cocaine addiction, it has remained unclear how risk-seeking behaviours observed in chronic cocaine users (CU) come about. Here we therefore test whether risky decision-making observed in CU is driven by alterations in individual sensitivity to the available information (*gain*, *loss*, and *risk*).

**Method:** A sample of 96 participants (56 CU and 40 controls) performed the no-feedback (“cold”) version of the Columbia Card Task. Structured psychiatric interviews and a comprehensive neuropsychological test battery were additionally conducted. Current and recent substance use was objectively assessed by toxicological urine and hair analysis.

**Results:** Compared to controls, CU showed increased risk-seeking in unfavourable decision scenarios in which the risk was high and the returns were low, and a tendency for increased risk aversion in favourable decision scenarios. These differences arose from the fact that CU were less sensitive to *gain*, but similarly sensitive to *loss* and *risk* information in comparison to controls. Further analysis revealed that individual differences in sensitivity to *loss* and *risk* were related to cognitive performance and impulsivity.

**Conclusion:** The reduced sensitivity to *gain* information in people with CU may contribute to their propensity for making risky decisions. While these alterations in the sensitivity to *gain* might be directly related to cocaine use per se, the individual psychopathological profile of CU might moderate their sensitivity to *risk* and *loss* impulsivity.

## 1. Background

Goal-oriented behaviour enables survival and can be viewed as the product of value-based decisions that relate the returns (i.e., the gains minus the losses) to the risks (e.g., uncertainty of returns or probability of a loss) of different courses of action in an individual-specific fashion [1–4]. Value-based decision processes can be affected by several factors, for instance, the degrees of uncertainty associated with the decision [5], development stages [5], social contexts [6], and several psychiatric disorders [7]. Concerning the to the last one, decision-making impairments often represent one of the main behavioural characteristics of substance-related disorders, contributing both to the impulsive initiation of substance use and to the compulsive maintenance of the addictive behaviour [8]. Such deficits seem to be even more severe when substances with strong addictive potentials are involved [9, 10], such as cocaine [11, 12].

Despite its negative consequences related with its use, cocaine remains one of the most commonly used illicit substances [13, 14]. In addition to the immediate risk of overdose and intoxication, cocaine use represents a substantial burden for the individual and their families as for the society because of its associations with cardiovascular [15], neurological [10, 16–18], and psychiatric [19] disorders, as well as with cognitive deficits [20, 21]. The negative consequences of cocaine use include not only decreases in quality of life and social functioning [3], but also an increase in high-risk behaviours and drug usage [11, 22]. From a clinical perspective, chronic cocaine use is usually accompanied by increased forgoing of occupational or recreational activities and by an increase of cocaine-seeking behaviours [23], which, from a decision-making viewpoint, suggest an alteration in the sensitivity of value-based decisions to the risks and returns of different courses of action.

Neurobehavioral research has sought to identify the neuroplastic adaptations underlying disadvantageous decision-making [23, 24], suggesting that chronic cocaine users (CU) are less sensitive to gains (i.e., magnitude of positive outcomes) and losses (i.e., magnitude of negative outcomes) in everyday situations [25, 26]. Particularly, CU have been proposed to suffer from a generalised impairment in value representation, reflected in blunted neural responses to non-substance-related (social and non-social) rewards, specifically in value-coding regions such as ventromedial prefrontal cortex [3, 27, 28]. Based on these findings we hypothesize that the deficits of chronic cocaine use in risky decision-making partially arise from alterations in return sensitivity.

Moreover, chronic cocaine use is associated with neuroplastic adaptations in brain networks involved in executive functioning and risk-taking, as indicated by reduced cortical thickness in the lateral prefrontal cortex, anterior cingulate cortex, and orbitofrontal cortex [20, 29, 30]. Such alterations may partially explain why CU are more likely to make maladaptive decisions in situations requiring implicit learning about risks, and why they prefer up-front high gain at the cost of higher risk [31–33]. Thus, it has also been hypothesized that CU may underestimate the risk of losing occurring over an extended period, resulting in long-term losses exceeding short-term gains; a phenomenon previously described as “myopia for the future” [34, 35]. Accordingly, we also hypothesize that CU are less sensitive to the risks of particular courses of action.

Together, although the literature suggests that CU may show impaired weighing or estimation of risks and returns in value-based decisions, so far, no study has investigated whether CU show decreased sensitivity particularly to information about gain magnitude, loss magnitude, and the probability of losing (i.e., risks). Moreover, it remained unclear whether such alteration can explain their risk-taking behaviour in decision scenarios with varying expected value. Here we aimed to fill these gaps, applying the no-feedback (“cold”) version of the Columbia Card Task, CCT [36] – a task designed to assess risky decision-making as a function of individual sensitivity to gains, losses, and risk. In addition, we also aimed to precisely investigate the effects of demographic, cognitive, psychopathological, and substance use severity on the sensitivity gain, loss, and risk information. Hence, based on previous studies that indicate that self-reported impulsivity and gambling behaviour are strongly state-dependent in CU [37], and that the often comorbid symptoms of an attention deficit hyperactivity disorder (ADHD) aggravates the effects of cocaine use on cognitive impairment [38, 39], we expect that both trait impulsivity and ADHD symptoms would reduce sensitivity to gain, loss, and risk information in CU. In sum, we investigate how chronic cocaine use, as well as demographic, clinical, and cognitive factors, affect sensitivity to gain, loss, and risk information in value-based decisions. Our findings will provide a basis for a better understanding of the proclivity of CU for risky behaviours. This knowledge may provide leads on how to improve the efficiency and the efficacy of preventive and therapeutic strategies.

## 2. Methods

### 2.1 Participants

In the context of the *Stress and Social Cognition Study* (SSCP), a total sample of 123 participants (69 chronic CU and 54 stimulant-naïve controls) was assessed (for detailed information on recruitment procedure, please see Supplementary Material). CU were included in the study if cocaine was the primary used illegal drug, if a lifetime cumulative consumption of at least 100g of cocaine was estimated by self-report and if current abstinence duration was <6 months. After applying the exclusion criteria (for detailed information on exclusion criteria, please see Supplementary Material), a total sample of 99 participants (59 chronic CU and 40 stimulant-naïve controls) was considered. However, two participants could not perform the CCT for technical reasons and one participant was excluded because the CCT data revealed random responses, suggesting that the participant did not understand the task or was not sufficiently motivated to perform the task. Therefore, 96 participants (56 chronic CU and 40 stimulant-naïve controls) matched for sex, age, smoking status, and weekly alcohol use (average number of times people drink per week) were analysed in this study. The study was approved by the *Cantonal Ethics Committee of Zurich* (BASEC ID 2016-00278) and preregistered at the *International Standard Randomised Controlled Trial Number* (ISRCTN-10690316). All participants provided written informed consent in accordance with the Declaration of Helsinki and were compensated for their participation.

### 2.2 Clinical and substance-related assessment

The psychopathological assessment was carried out with the Structured Clinical Interview I for the DSM-IV-R (SCID-I) [40]. ADHD symptoms were collected with the ADHD self-rating scale (ADHD-SR) [41]. Trait impulsivity was measured with the Barratt Impulsiveness Scale (BIS) [42]. Self-reported drug use was assessed with the structured and standardized Interview for Psychotropic Drug Consumption [43].

### 2.3 Urine and hair toxicological analysis

Urine analyses using a semiquantitative enzyme multiplied immunoassay method targeted the following substances: amphetamines, barbiturates, benzodiazepines, cocaine, methadone, morphine-related opiates, and tetrahydrocannabinol. In addition, quantitative analysis of hair samples using liquid chromatography tandem mass spectrometry (LC-MS/MS) was used to investigate substance consumption over the last 4 months as represented in the proximal 4cm-segment of the hair samples. In total 88 compounds were assessed. For a complete description of all compounds accessed, please see Supplementary Material.

### 2.4 General cognitive assessment

The German vocabulary test *Mehrfachwahl-Wortschatz-Intelligenztest* (MWT-B) was applied to estimate premorbid verbal intelligence [44]. General cognitive performance was assessed with a selection of three tasks from the *Cambridge Neuropsychological Test Automated Battery* (CANTAB, http://www.cantab.com): the *Spatial Working Memory* task (SWM) (to assess working memory and executive functioning), the *Match to Sample Visual Search* task (MTS) (a visual matching test involving a trade-off of speed and accuracy), and the *Rapid Visual Information Processing* task (RVP) (to assess sustained attention capacity). For detailed information about these tasks, please see the Supplementary Material.

### 2.5 Columbia Card Task

Due to our primary focus on understanding how the sensitivity to *gain*, *loss*, and *risk* information can explain people’s behaviour in different decision scenarios, participants performed the no-feedback condition of the CCT. In the CCT participants faced a deck with 32 facedown cards and three explicit pieces of information (i.e., scenario properties): how many losing cards were hidden in the deck (i.e., *risk*, 1 or 3), the amount associated with each losing card (i.e., *loss*, −250 or −750 points) and the amount associated with each winning card (i.e., *gain*, 10 or 30 points). In every round, participants decided how many cards the computer would randomly select and turn over, knowing that the round would end immediately if the computer selected one of the losing cards. The different combinations of *gain*, *loss*, and *risk* culminated in eight possible decision scenarios that can be sorted from the most favourable to the least favourable, according to the expected value. For additional information about the CCT, please see the Supplementary Material.

#### 2.5.1. Risk-attitude

The primary outcome of the CCT is the average number of cards chosen, which can be interpreted as a general proxy of risk-seeking behaviour, with a higher number of cards corresponding to greater risk-proneness [36, 45–47]. We also analysed the risk-seeking behaviour separately for each decision scenarios in order to assess risk-taking in a more fine-grained fashion.

#### 2.5.2. Sensitivity to gain, loss, and risk

Concerning sensitivity to the scenario properties (i.e., *gain*, *loss*, and *risk*), a normative analysis of the CCT suggests that participants should choose the number of cards in accordance with their belief that the subjective value of that number of cards is maximal [36], in which an optimal strategy take into account *gain*, *loss*, and *risk*. It can be analysed at both the group and individual level [36].

At the group level we performed a linear mixed effect model (LMM) [48] including group (CU or stimulant-naïve controls), *gain* (10, 30), *loss* (−250, −750), and *risk* (1, 3) as fixed-effects. This model allowed us to extract the estimates of regression coefficients for both group and the scenario properties. The LMM accounted for the random-effects of each participant slope and intercept associated with the different scenario properties [48]. Because the estimates of the regression coefficients represent the slope of the function (i.e., the weighting that *gain*, *loss*, and *risk* received in determining the number of cards), we used these values as measures for the sensitivity to *gain, loss*, and *risk*.

At the individual level, an LMM analyses were performed for each participant separately to investigate how the sensitivity to *gain*, *loss*, and *risk* influenced his/her risk-taking. Random intercepts for the three blocks and the 24 rounds were included in the model. The number of cards chosen was mean centred according to the control group. Similarly, to the group analysis, three coefficients were extracted for each participant, each one capturing how the participant weighted the *gain*, *loss*, and *risk* factors.

### 2.6 Statistical analysis

All statistical analyses were performed with the open source statistical software R [49]. Regarding demographic, clinical, cognitive, and substance-related variables, frequency data were analysed by means of Pearson’s chi-square tests and quantitative data by Student’s t-tests or, when data was non-normally distributed, Wilcoxon rank sum tests (i.e., Shapiro–Wilk W < .001, and skew and kurtosis divided by 2 standard errors < 2).

To assess potential group differences in overall risk-attitude, independent of the decision scenario, we used several LMM including different random intercepts and slopes and tested them with the model fitting function “anova” [50]. The best model included a random slope and intercept for each participant and scenario properties (i.e., *gain*, *loss*, and *risk*). Then, using a similar strategy of testing different random intercepts and slopes with the same model fitting function, the effect of group on the average number of cards chosen at each decision scenario separately was investigated including a random intercept for participant only.

Secondly, to investigate group differences in the sensitivity to scenario properties, we first performed an LMM analysis including group (CU or stimulant-naïve controls) and the expected value of each decision scenario as fixed-effects, and a random slope and intercept for each participant and the three scenario properties (*gain*, *loss*, and *risk*). Then, as previously mentioned, an LMM analysis including group, *gain*, *loss*, and *risk* as fixed-effects and a random slope and intercept for each participant and the three scenario properties was performed. To explore within group variance explained by the use of information, the same model was also analysed for both groups separately. Effect sizes were calculated by Pearson’s correlation coefficient (0≤|r|<.10 small effect size; .10≤|r|<.30 medium effect size; .30≤|r|<.50 large effect size), which has been suggested as a versatile measure of the strength of an experimental effect with an intuitive interpretation – absolute values of “r” are constrained to lie between 0 (no effect) and 1 (maximal effect) [51].

In a third step, we examined whether the reported demographic, cognitive and clinical group differences contribute to the sensitivity to *gain, loss*, and *risk* that was extracted for each participant individually through LMM. To do so, we performed hierarchical linear models including years of education, verbal IQ, SWM Strategy score, SWM Total error score, meta-efficiency index, trait impulsivity, and ADHD symptoms. To test for multicollinearity between predictors we performed a set of Spearman’s rank correlations over all participants. Based on the cut-offs suggested by Cohen [51], predictors with large effect size correlations were not included together in the same model. Then, to identify the subset of variables with the highest explanatory power we incorporated the predictors into the model ‘one by one’. As before, models were compared using the model fitting function “anova” [50].

Subsequently, linear regressions were performed within both groups to relate individual sensitivity to *gain*, *loss*, and *risk* to the average number of cards chosen in each decision scenario controlling for the predictors in which a significant effect was found in the hierarchical linear models. Finally, to analyse use severity, we ln10-transformed hair metabolites measures (due to the highly right-skewed distribution and the resulting deviation from the normal distribution). We then conducted linear regression analyses within CU to examine how substance-related variables related to *gain, loss*, and *risk* sensitivity, controlling for the predictors in which a significant effect was found in the hierarchical linear models.

## 3. Results

### 3.1 Demographic characteristics and substance use

As intended by our matching procedure, the groups did not differ regarding age, sex as well as nicotine and cannabis smoking status (Table 1 and Table 2), although on average CU had fewer years of education than stimulant-naïve controls and lower verbal IQ. As expected, CU displayed higher ADHD-SR scores and higher trait impulsivity in the BIS. Cognitive assessment revealed that CU exhibited worse working memory and executive functioning, measured by the SWM between/total errors and SWM strategy score, respectively. CU also showed lower signal detection/sustained attention in the RVP and lower efficiency indices in the RVP and MTS.

**Table 1.** Demographic, cognitive and clinical data.

|  | Controls (n=40) | Cocaine Users (n=56) | Test Statistics | df | *p* |
| --- | --- | --- | --- | --- | --- |
| *Demographics* |  |  |  |  |  |
| Age, y | 29.3 (7.1) | 32.3 (7.9) | *t* = -1.9 | 89.0 | .060 |
| Sex, f/m | 17 / 23 | 17 / 39 | $\chi^2 = 1.5$ | 1 | .220 |
| Verbal IQ <sup>a</sup> | 100.6 (6.5) | 95.4 (5.8) | *t* = 3.9 | 77.7 | **.000** |
| School education, y | 10.2 (1.4) | 9.4 (.89) | *W* = 1415 | - | **.009** |
| *Cognition* |  |  |  |  |  |
| SWM - Between errors | 15.9 (13.7) | 26.0 (17.3) | *W* = 723 | - | **.003** |
| SWM - Within errors | .85 (2.3) | 1.1 (2.1) | *W* = 952 | - | .151 |
| SWM - Total errors | 16.3 (14.1) | 26.4 (17.4) | *W* = 729 | - | **.003** |
| SWM - Strategy score | 29.0 (6.4) | 32.5 (5.8) | *t* = -2.7 | 78.9 | **.007** |
| RVP - Response A' | .92 (.06) | .88 (.05) | *W* = 1638 | - | **.000** |
| RVP - Response bias B' | .89 (.32) | .93 (.10) | *W* = 1121 | - | .655 |
| RVP - Mean latency, ms. | 405.9 (196.1) | 425.4 (91.4) | *t* = -.94 | 76.1 | .350 |
| RVP - Total false alarms | 1.2 (1.4) | 2.3 (5.2) | *W* = 835 | - | **.028** |
| RVP - Impulsivity Index | .00 (.93) | .66 (3.5) | *W* = 1005 | - | .392 |
| RVP - Efficiency Index | .00 (1.7) | -1.0 (4.1) | *W* = 1477 | - | **.007** |
| MTS - Correct, % | 96.1 (5.0) | 93.8 (6.8) | *W* = 1306 | - | .097 |
| MTS - Correct reaction time, ms. | 2568 (623.9) | 2662 (737.3) | *t* = -.67 | 90.8 | .504 |
| MTS - Time change 2-8, ms. | 146.0 (303.5) | 139.2 (434.2) | *W* = 1193 | - | .480 |
| MTS - Impulsivity Index | .00 (1.5) | -.29 (3.1) | *t* = .61 | 84.3 | .543 |
| MTS - Efficiency Index | .00 (1.3) | -1.1 (2.9) | *t* = 2.55 | 80.2 | **.012** |
| *BIS subscales* |  |  |  |  |  |
| Total Score | 64.2 11.1 | 70.0 12.0 | *t* = -2.4 | 87.8 | **.016** |
| Attention Impulsiveness | 14.6 (3.8) | 16.9 (4.4) | *t* = -2.7 | 90.6 | **.007** |
| Motor Impulsiveness | 23.4 (5.1) | 24.4 (5.2) | *t* = -.94 | 85.0 | .346 |
| Non-planning Impulsiveness | 26.1 (4.8) | 28.6 (4.9) | *t* = -2.4 | 84.8 | **.014** |
| *Clinical* |  |  |  |  |  |
| ADHD, y/n <sup>b</sup> | 7/33 | 43/13 | $\chi^2 = .46$ | 1 | .496 |
| ADHD sum score | 10.5 (9.7) | 14.6 (10.1) | *W* = 827 | - | **.029** |

Note. Table reports counts or means with standard deviations in brackets. Significant group differences are shown in bold. *t* = Student *t*-test; $\chi^2$ = Pearson chi-square; *W* = Wilcoxon rank sum test. (a) Verbal intelligence quotient estimated by the Mehrfachwahl-Wortschatz-Intelligenztest (MWT-B); (b) Cut-off DSM-IV criteria as assessed by the ADHD-SR questionnaire. RVP, Rapid Visual Information Processing task; SWM, Spatial Working Memory task; MTS, Match to Sample Visual Search task. BIS, Barratt Impulsiveness Scale. ADHD, Attention deficit hyperactivity disorder.

**Table 2.** Substance use related disorders and drug consumption pattern.

|  | Controls (n=40) | Cocaine Users (n=56) | Test Statistics | df | p |
| --- | --- | --- | --- | --- | --- |
| *Nicotine* |  |  |  |  |  |
| Smoking, y/n | 37/3 | 53/3 | $\chi^2 = .18$ | 1 | .668 |
| Cigarettes per week <sup>a</sup> | 67.0 (47.1) | 111.6 (68.2) | W = 575.5 | - | **.000** |
| Years of use | 11.7 (6.1) | 17.2 (16.1) | W = 710 | - | **.026** |
| *Alcohol* |  |  |  |  |  |
| Times per week <sup>a</sup> | 2.7 (2.2) | 3.3 (3.5) | W = 1052 | - | .719 |
| Grams per week <sup>a, b</sup> | 22796 (24922) | 59247 (117708) | W = 972 | - | .334 |
| Years of use | 14.1 (6.9) | 16.8 (7.0) | t = -1.88 | 84.81 | .062 |
| Estimated cumulative lifetime dose, g <sup>b</sup> | 65609 (51775) | 284749 (428576) | W = 707 | - | **.002** |
| *Cocaine* |  |  |  |  |  |
| Cocaine lifetime experience, y/n <sup>c</sup> | 5/35 | 56/0 | - | - | - |
| Times per week <sup>a</sup> | - | 2.4 (2.4) | - | - | - |
| Grams per week <sup>a</sup> | - | 3.9 (5.76) | - | - | - |
| Years of use | - | 12.0 (7.5) | - | - | - |
| Estimated cumulative lifetime dose, g | - | 1919 (2290) | - | - | - |
| Abstinence period, days | - | 14.3 (23.7) | - | - | - |
| Cocaine, ng/mg in hair [n] <sup>d</sup> | - | 19388 (26967) [56] | - | - | - |
| Benzoylcegonine, ng/mg in hair [n] <sup>d</sup> | - | 11197 (15907) [56] | - | - | - |
| Cocathylene, ng/mg in hair [n] <sup>d</sup> | - | 960.8 (1943) [50] | - | - | - |
| Norcocaine, ng/mg in hair [n] <sup>d</sup> | - | 447.8 (654.5) [56] | - | - | - |
| Cocaine<sub>total</sub>, ng/mg in hair [n] <sup>d, e</sup> | - | 31034 (41770) [56] | - | - | - |
| Urine toxicology, n/p in hair [n] <sup>f</sup> | 40/0 | 33/22 | $\chi^2 = 33.5$ | 2 | **.000** |
| *Cannabis* |  |  |  |  |  |
| Cannabis lifetime experience, y/n <sup>c</sup> | 35/5 | 50/6 | $\chi^2 = .07$ | 1 | .395 |
| Grams per week <sup>a</sup> | .86 (1.9) | 2.0 (7.0) | W = 798 | - | .493 |
| Years of use | 7.8 (6.0) | 12.7 (9.0) | W = 594 | - | **.012** |
| Estimated cumulative lifetime dose, g | 142.8 (370.0) | 3341 (5608) | W = 385 | - | **.000** |
| Abstinence period, days | 1159.0 (2222.1) | 1005 (2216) | W = 1012 | - | .221 |
| THC, ng/mg in hair [n] | 36.4 (43.5) [5] | 137.0 (258.9) [16] | W = 26.5 | - | .264 |
| CBD, ng/mg in hair [n] | 16.0 (-) [1] | 32.1 (35.8) [8] | W = 4 | - | 1 |
| CBN, ng/mg in hair [n] | 16.6 (18.7) [3] | 54.5 (74.5) [14] | W = 14 | - | .376 |
| Urine toxicology, n/p <sup>f</sup> | 38/2 | 46/10 | $\chi^2 = 5.4$ | - | .066 |
| *MDMA* |  |  |  |  |  |
| MDMA lifetime experience, y/n <sup>c</sup> | 9/31 | 49/7 | $\chi^2 = 41.2$ | 1 | **.000** |
| Grams per week <sup>a</sup> | .00 (-) [1] | .04 (.08) | W = 131 | - | .054 |
| Years of use | 1.8 (1.6) | 7.8 (7.6) | W = 108 | - | **.015** |
| Estimated cumulative lifetime dose, g | .34 (.46) | 50.4 (154.1) | W = 40.5 | - | **.000** |
| Abstinence period, days | 1771 (3033) | 993.5 (1965.9) | W = 233.5 | - | .780 |
| MDMA, ng/mg in hair [n] <sup>d</sup> | 109.0 (55.5) [4] | 2579 (4976) [36] | W = 38.5 | - | .130 |
| MDA, ng/mg in hair [n] <sup>d</sup> | 6.6 (3.0) [3] | 189.2 (436.4) [29] | W = 19.5 | - | .120 |
| *Amphetamine* |  |  |  |  |  |
| Amphetamine lifetime experience, y/n <sup>c</sup> | 4/36 | 42/14 | $\chi^2 = 39.5$ | 1 | **.000** |
| Grams per week <sup>a</sup> | .01 (.02) | .17 (.63) | W = 54 | - | .241 |
| Years of use | .04 (.08) | 7.4 (6.5) | W = 7.5 | - | **.002** |
| Estimated cumulative lifetime dose, g | 2.1 (3.8) | 162.8 (369.8) | W = 39 | - | .079 |
| Abstinence period, days | 1771 (3033) | 993.5 (1965.9) | W = 233.5 | - | .780 |
| Amphetamine, ng/mg in hair [n] <sup>d</sup> | - (-) [0] | 1222 (1700) [13] | - | - | - |
| *SCID-I diagnosis* |  |  |  |  |  |
| Alcohol dependency current, y/n | 0/40 | 8/48 | $\chi^2 = 6.2$ | 1 | **.012** |
| Alcohol dependency past, y/n | 0/40 | 18/38 | $\chi^2 = 15.8$ | 1 | **.000** |
| Alcohol abuse current, y/n | 1/39 | 18/38 | $\chi^2 = 12.9$ | 1 | **.000** |
| Alcohol abuse past, y/n | 4/36 | 26/30 | $\chi^2 = 14.4$ | 1 | **.000** |
| Cocaine dependency current, y/n | 0/40 | 36/20 | $\chi^2 = 41.1$ | 1 | **.000** |
| Cocaine dependency past, y/n | 0/40 | 39/17 | $\chi^2 = 46.9$ | 1 | **.000** |
| Cocaine abuse current, y/n | 0/40 | 36/20 | $\chi^2 = 41.1$ | 1 | **.000** |
| Cocaine abuse past, y/n | 0/40 | 36/20 | $\chi^2 = 41.1$ | 1 | **.000** |
| Cannabis dependency current, y/n | 0/40 | 1/55 | $\chi^2 = .72$ | 1 | .395 |
| Cannabis dependency past, y/n | 0/40 | 9/46 | $\chi^2 = 7.2$ | 1 | **.007** |
| Cannabis abuse current, y/n | 0/40 | 4/52 | $\chi^2 = 2.9$ | 1 | .084 |
| Cannabis abuse past, y/n | 5/35 | 17/38 | $\chi^2 = 4.4$ | 1 | **.035** |

Note. Table reports counts or means with standard deviations in brackets. Significant differences are shown in bold. t = Student t-test; $\chi^2$ = Pearson chi-square; W = Wilcoxon rank sum test. Here we specifically reported the most prevalent substances and metabolites: THC, Tetrahydrocannabinol; CBD, cannabinoide; CBN, cannabinoide; MDMA, 3,4-Methylenedioxymethamphetamine; MDA, 3,4-Methylenedioxymphetamine; SCID-I, Structured Clinical Interview for DSM-IV Axis I Disorders. (a) Average use of the current consumption period. (b) Pure alcohol estimation; (c) Self-report: Have you ever consumed this substance, at least once, in your life? (d) Cut-off values for cocaine = 500 pg/mg and for amphetamines/MDMA = 200 pg/mg (Cooper et al., 2012). (e) Cocaine<sub>total</sub> (= Cocaine + Benzoylcegonine + Norcocaine) is a more robust procedure for discrimination between incorporation and contamination of hairs (Hoelzle et al., 2008). (f) Urine toxicology (neg/pos) are based on cut-off value for Cocaine = 150 ng/ml and for Tetrahydrocannabinol 50 ng/ml (Substance Abuse and Mental Health Services Administration, 2008).

Hair samples revealed a clear dominance of cocaine compared with all other illegal drugs, as set out by the inclusion criteria (in the mean 12 times more cocaine than MDMA and 25 times more cocaine than amphetamines) (Table 2). SCID-I revealed a higher frequency of alcohol and cannabis-related disorders in the CU compared to stimulant-naïve controls. Pearson correlations revealed that total hair concentration of cocaine metabolites (Cocaine_total_) correlated with self-reported estimated cumulative dose (r=.366, *p*<.01, n=56), duration of use (r=.326, *p*<.05, n=56), and days of abstinence before the measurement (r=-.333, *p*<.05, n=56).

### 3.2 Decision-making

#### 3.2.1. Overall and scenario-specific risk-attitude

To investigate group differences in overall risk-attitude, we performed a LMM including group as a predictor and a random slope for each participant at each scenario property. The analysis revealed that CU (Mean = 12.37, SD = 8.1) did not differ from stimulant-naïve controls (Mean = 11.67, SD = 8.5) concerning the average number of cards chosen over all decision scenarios (ß = .008, 95%CI = −1.84 to 1.86, t[94] = .008, *p* = .993, r = .0009).

Afterwards, we investigated risk-taking for each scenario independently by modelling a different random intercept for each participant. We found that CU chose more cards than stimulant-naïve controls in high risk, low return scenarios (for the most unfavourable decision scenario: ß = 3.67, 95%CI = 1.05 to 6.30, t[94] = 2.76, *p* = .006; Figure 1). This finding remained after including verbal IQ, years of education, and ADHD symptoms as covariates (ß = 3.50, 95%CI = 3.98 to 4.86, t[91] = 2.31, *p* = .022; Figure 1). Additionally, we found that CU tended to choose fewer cards than stimulant-naïve controls in low risk, high return scenarios (for the most favourable decision scenario: ß = −2.56, 95%CI = −5.27 to .14, t[94] = −1.87, *p* = .064; Figure 1), although this finding did not reach significance. Thus, CU were more risk-taking than controls in unfavourable decision scenarios but tended to decide more cautiously in favourable decision scenarios.

**Figure 1.**
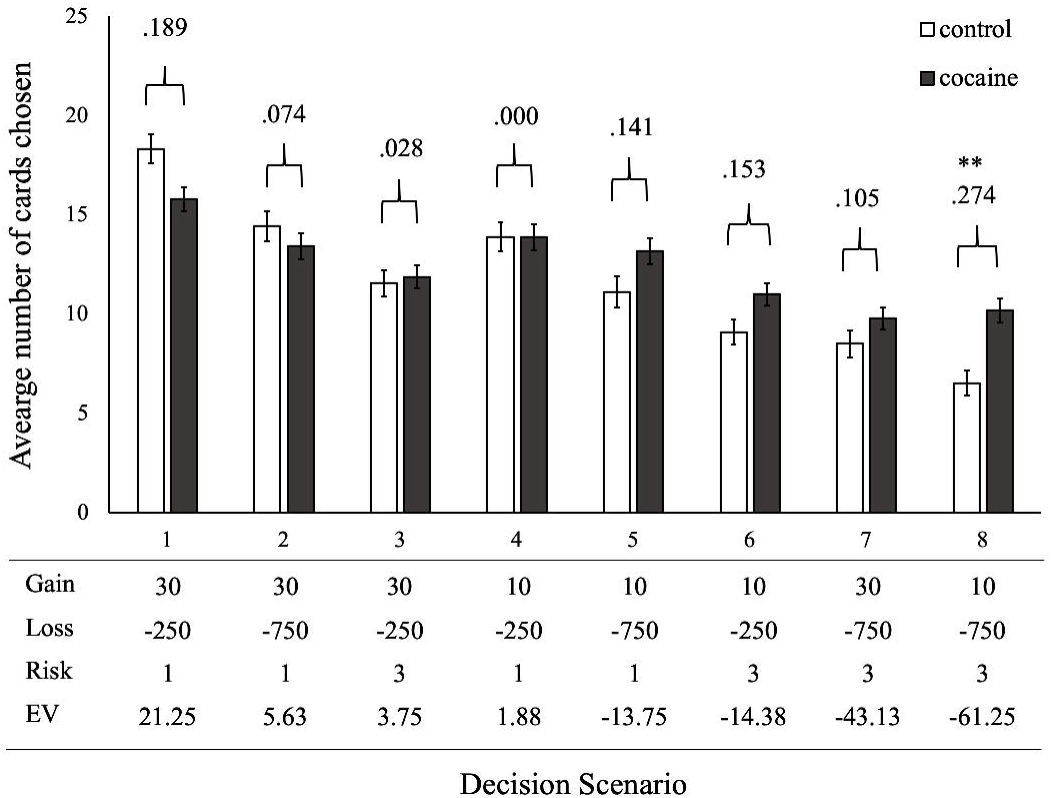
Average number of cards selected in each scenario. EV, Expected Value. Effect sizes were calculated by Pearson’s correlation coefficient (r <.10 small effect size; r <.30 medium effect size; r <.50 large effect size). ** *p*-value <.01.

#### 3.2.2. Sensitivity to gain, loss, and risk

To investigate group differences in overall sensitivity to the expected value, an LMM analysis including group and expected value as fixed effects, as well as random slopes and intercepts for each participant and each scenario property was performed. The data revealed that CU are significantly less sensitive to the expected value (ß = -.052, 95%CI = -.08 to -.02, t[2206] = −3.55, *p* = .0004, r = .075) than stimulant-naïve controls. Subsequently, to investigate group differences in the use of scenario properties, we performed an LMM analysis including group, *gain*, *loss*, and *risk* as fixed-effects and random slopes and intercepts for each participant and each scenario property. As shown in Figure 2, we found a significant interaction of group with gain and a marginally significant interaction of group with *loss*. These interactions suggest that when the gain is high and, to a lesser degree, when the loss is low, CU select fewer cards than stimulant-naïve controls. We found no interaction of group with risk. Moreover, as expected from the preceding analysis (section 3.2.1), main effects were found for *gain*, *loss*, and *risk* but not for group. Confirming our hypothesis, these findings suggest that, compared to controls, CU are less sensitive to the expected value (i.e., the “favourableness” of the decision scenarios). In particularly, CU are less sensitive to *gain* information, choosing fewer cards than controls at high gains.

**Figure 2.**
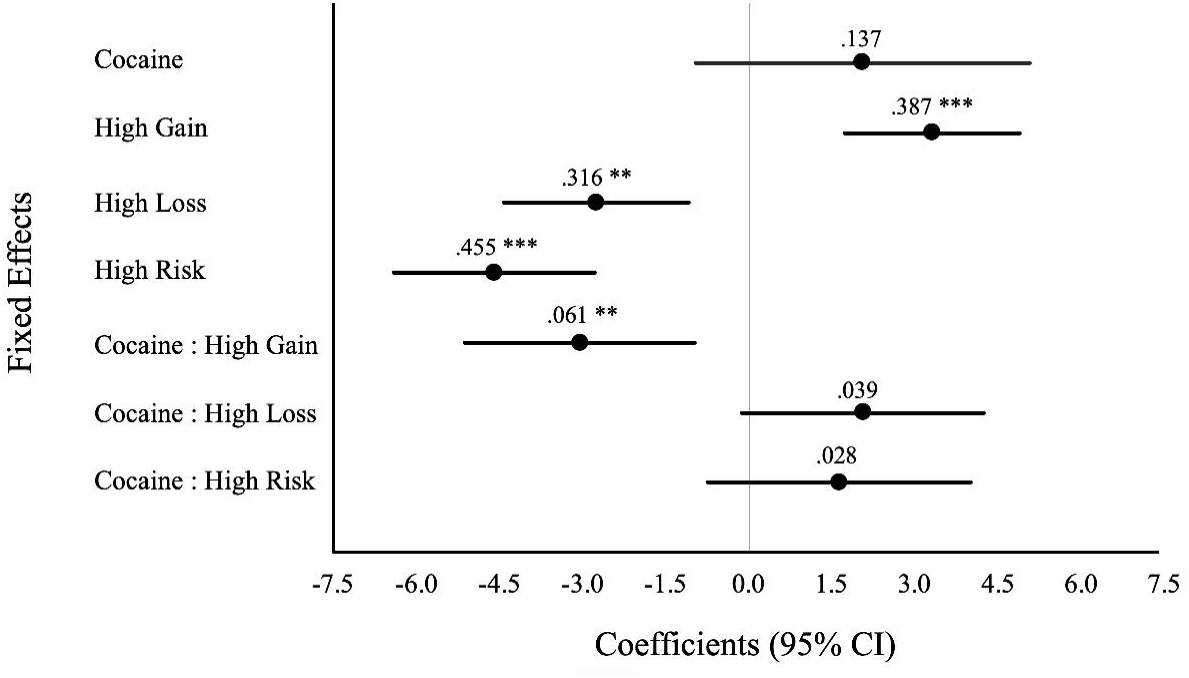
Coefficient estimates for the main fixed effects and interactions of the linear mixed model. The model included random slopes and intercepts for each participant and each scenario property (i.e. gain, loss and risk). Stimulant-naïve control group, at low gain, low loss and low risk was used as reference group. Of note, CU were less sensitive to gain than control participants. Conditional R^2^ = .54, marginal R^2^ = .10. CU, chronic cocaine users. *** *p*-value <.001; ** *p*-value <.01.

To explore within-group variance explained by the use of scenario properties, we analysed CU and stimulant-naïve controls separately. The control group displayed a significant effect of all scenario properties (*gain*: ß = 3.30, 95%CI = 1.70 to 4.91, t[913] = 4.03, *p* < .001, r = .132; *loss*: ß = 2.76, 95%CI = .99 to 4.54, t[913] = 3.04, *p* = .002, r = .100; *risk*: ß = −4.60, 95%CI = −6.47 to −2.74, t[913] = −4.82, *p* < .001, r = .157) on the number of cards chosen, implying that controls chose more cards when the gain was high and loss or risk were low (condition R^2^ = .59; marginal R^2^ = .16). Within the CU group we found a significant effect for *risk* (ß = −2.97, 95%CI = −4.48 to −1.47, t[1281] = −3.86, *p* = .000, r = .107) but not for *gain* (ß = .25, 95%CI = −1.08 to 1.58, t[1281] = .36, *p* = .714, r = .010) or *loss* (ß = -.71, 95%CI = -.63 to 2.06, t[1281] = −1.03, *p* = .299, r = .028; condition R^2^ = .50; marginal R^2^ = .05). Together, these results suggest that CU were predominantly sensitive to *risk*, while controls were sensitive also to *gain* and *loss* information.

#### 3.2.3. Impact of demographic, cognitive, and clinical variables

To examine whether the reported group differences (Table 1) relate to differential weighing of *gain, loss, and risk* information, we used hierarchical linear models. Since the SWM Strategy score and the SWM Total error score revealed a large effect size correlation, as well as the BIS total score and the ADHD sum score, these variables were entered into separate models (see Supplementary Table S1). As shown in Table 3, *gain* and *loss* sensitivity were best explained by a model that included group and years of school education (F[93]=8.36; R^2^=.134; *p*=.055; and F[93]=4.12; R^2^=.61; *p*=.054, respectively), suggesting that longer education leads to higher sensitivity to gains and losses. With regard to *risk* sensitivity, we found that the model with group, IQ, SWM Strategy score, and ADHD symptoms explained more variance than the other models (Table 3) (F[91]=5.40; R^2^=.156; *p*=.053). Additional multiple regressions did not reveal any effect for sex and age. Together, these data suggest that while gain sensitivity was explained primarily by group (and years of school education), loss and risk sensitivity were better explained by additional demographic, cognitive and clinical variables.

**Table 3.** Hierarchical multiple linear regression models for gain, loss and risk sensitivity..

|  | Cocaine | Years of Education | IQ | SWM Strategy | SWM Total Errors | Meta-Efficiency Index | BIS | ADHD | [df] F | R<sup>2</sup> | Model Comparison | *p* |
| --- | --- | --- | --- | --- | --- | --- | --- | --- | --- | --- | --- | --- |
| *Gain* |  |  |  |  |  |  |  |  |  |  |  |  |
| Model 1 | -.11 (-3.54) *** |  |  |  |  |  |  |  | [94] 12.59 | .109 | 1 vs. Null | **.000** |
| Model 2 | -.09 (-2.76) ** | .02 (1.94) * |  |  |  |  |  |  | [93] 8.36 | .134 | 2 vs. 1 | **.055** |
| Model 3 | -.11 (-3.01) ** | .03 (2.14) * | -.00 (-1.22) |  |  |  |  |  | [92] 6.09 | .138 | 3 vs. 2 | .225 |
| Model 4 | -.09 (-2.57) * | .02 (1.59) |  | -.00 (-.75) |  |  |  |  | [92] 5.74 | .130 | 4 vs. 2 | .449 |
| Model 5 | -.09 (-2.65) ** | .02 (1.85) |  |  | -.00 (-.11) |  |  |  | [92] 5.51 | .124 | 5 vs. 2 | .909 |
| Model 6 | -.08 (-2.45) * | .02 (1.90) |  |  |  | .00 (.1.01) |  |  | [92] 5.92 | .134 | 6 vs. 2 | .311 |
| Model 7 | -.10 (-2.81) ** | .02 (2) * |  |  |  |  | .00 (.60) |  | [92] 5.65 | .128 | 7 vs. 2 | .545 |
| Model 8 | -.09 (-2.64) ** | .02 (1.91) |  |  |  |  |  | -.00 (-.36) | [92] 5.56 | .126 | 8 vs. 2 | .716 |
| Full model <sup>a, b</sup> | -.10 (-2.69) ** | .02 (1.87) | -.00 (-1.21) | -.00 (-.60) | - | -.00 (-.62) | .00 (.62) | - | [89] 3.26 | .124 | Full vs. 2 | .558 |
| *Loss* |  |  |  |  |  |  |  |  |  |  |  |  |
| Model 1 | -.00 (-2.08) * |  |  |  |  |  |  |  | [94] 4.33 | .033 | 1 vs. Null | **.040** |
| Model 2 | -.00 (-1.35) | .00 (1.94) * |  |  |  |  |  |  | [93] 4.12 | .061 | 2 vs. 1 | **.054** |
| Model 3 | -.00 (-1.19) | .00 (1.85) | .00 (.24) |  |  |  |  |  | [92] 2.74 | .052 | 3 vs. 2 | .806 |
| Model 4 | -.00 (-1.03) | .00 (1.30) |  | -.00 (-1.78) |  |  |  |  | [92] 3.88 | .083 | 4 vs. 2 | .076 |
| Model 5 | -.00 (-.99) | .00 (1.57) |  |  | -.00 (-1.50) |  |  |  | [92] 3.54 | .074 | 5 vs. 2 | .135 |
| Model 6 | -.00 (-1.10) | .00 (1.91) |  |  |  | .00 (.94) |  |  | [92] 3.04 | .060 | 6 vs. 2 | .349 |
| Model 7 | -.00 (-1.45) | .00 (2.01) * |  |  |  |  | .00 (.67) |  | [92] 2.88 | .056 | 7 vs. 2 | .501 |
| Model 8 | -.00 (-1.47) | .00 (1.97) * |  |  |  |  |  | .00 (.83) | [92] 2.97 | .058 | 8 vs. 2 | .406 |
| Full model <sup>a, c</sup> | -.00 (-.95) | .00 (1.27) | .00 (.19) | -.00 (-1.64) | - | .00 (.42) | - | .00 (.99) | [89] 2.10 | .065 | Full vs. 2 | .364 |
| *Risk* |  |  |  |  |  |  |  |  |  |  |  |  |
| Model 1 | 1.08 (2.41) * |  |  |  |  |  |  |  | [94] 5.84 | .017 | 1 vs. Null | **.017** |
| Model 2 | .97 (2.04) * | -.13 (-.70) |  |  |  |  |  |  | [93] 3.15 | .043 | 2 vs. 1 | .480 |
| Model 3 | .67 (1.42) |  | -.07 (-2.21) * |  |  |  |  |  | [93] 5.50 | .086 | 3 vs. 1 | **.029** |
| Model 4a | .42 (.88) |  | -.07 (-2.02) * | .08 (2.38) * |  |  |  |  | [92] 5.74 | .130 | 4a vs. 3 | **.019** |
| Model 4b <sup>a</sup> | .41 (.84) |  | -.07 (-2.21) * | - | .02 (2.04) * |  |  |  | [92] 5.19 | .116 | 4b vs. 4a | - |
| Model 5 | .40 (.82) |  | -.07 (-2.02) * | .08 (2.20) * |  | -.02 (-.21) |  |  | [91] 4.27 | .121 | 5 vs. 4a | .828 |
| Model 6 | .62 (1.30) |  | -.06 (-1.97) * | .09 (2.56) * |  |  | -.03 (-1.96) * |  | [91] 5.40 | .156 | 7 vs. 4a | **.053** |
| Model 7 <sup>c</sup> | .61 (1.31) |  | -.07 (-2.05) * | .09 (2.71) ** |  |  | - | -.05 (-2.58) * | [91] 6.25 | .181 | 7 vs. 6 | - |
| Full model <sup>a, c</sup> | .64 (1.30) | .07 (.38) | -.07 (-2.07) * | .09 (2.53) * | - | -.01 (-.10) | - | -.05 (-2.54) * | [88] 4.11 | .164 | Full vs. 8 | .921 |

Note. IQ, Intelligence Quotient assessed with the Mehrfachwahl-Wortschatz-Intelligenztest (MWT-B); SWM, Spatial Working Memory task; BIS, total sum score from Barratt Impulsiveness Scale; ADHD, Attention deficit hyperactivity disorder total sum score from ADHD self-rating scale. *p*-value $\leq .05$ \*; *p*-value $< .01$ \*\*; *p*-value $< .001$ \*\*\*. The table reports adjusted R<sup>2</sup>. Bold *p*-values indicate models with more favourable fit. (a) Due to multicollinearity, EF was not included in the same model as WM. (b) Due to multicollinearity, BIS total score but not ADHD was included in the model. (c) Due to multicollinearity, ADHD total sum score but not BIS was included in the model. (d) Efficiency Index is the average of the efficiency indices from the Rapid Visual Information Processing task and the Match to Sample Visual Search efficiency task.

Next, we aimed to investigate how *gain, loss, and risk* sensitivity correlate with risk-attitude in each decision scenario, after correcting for the significant effects found in the best explanatory hierarchical linear models (Table 4). Specifically, *gain* and *loss* correlations were corrected for years of school education and *risk* correlations were corrected for IQ, executive functioning, and ADHD symptoms. Our data revealed that, within the control group, in the most and the least favourable decision scenario, risk-attitude correlated with the sensitivity to *gain*, *loss* and *risk* information. However, within the CU group, only sensitivity to *gain*, *loss* correlated with risk-attitude in the least favourable decision scenario, while only the sensitivity to *risk* information correlated with risk-attitude in the most favourable decision scenario.

**Table 4.** Correlations between gain, loss and risk sensitivity and risk-attitude at each decision scenario.

|  | Non-Stimulant Controls |  |  | Cocaine Users |  |  |
| --- | --- | --- | --- | --- | --- | --- |
|  | Gain <sup>a</sup> | Loss <sup>a</sup> | Risk <sup>b</sup> | Gain <sup>a</sup> | Loss <sup>a</sup> | Risk <sup>b</sup> |
| *Average n° of cards* |  |  |  |  |  |  |
| Scenario 1 | .538 *** | .559 *** | -.583 *** | .193 | .184 | -.554 *** |
| Scenario 2 | -.135 | -.239 | -.503 ** | -.416 ** | .012 | -.472 *** |
| Scenario 3 | -.011 | -.394 * | -.144 | -.157 | -.320 * | -.485 *** |
| Scenario 4 | .456 ** | -.574 *** | .099 | -.406 ** | -.455 *** | -.357 ** |
| Scenario 5 | .088 | -.217 | .258 | .025 | .203 | .094 |
| Scenario 6 | -.530 *** | -.402 * | .417 * | -.557 *** | -.067 | .125 |
| Scenario 7 | -.244 | -.591 *** | .465 ** | -.057 | -.510 *** | .060 |
| Scenario 8 | -.439 ** | -.743 *** | .469 ** | -.523 *** | -.338 * | .120 |

Note. $p$ -value < .05 \*; $p$ -value < .01 \*\*; $p$ -value < .001 \*\*\*. (a) Corrected for years of school education; (b) Corrected for IQ, executive functioning and ADHD symptoms;

#### 3.2.3. Impact of cocaine use severity

Finally, we investigated how *gain, loss, and risk* sensitivity relate with cocaine-related metabolites and cocaine consumption self-report. Although no effect was found for the self-report estimated cumulative lifetime dose, we found a negative effect of level of benzoylecgonine (r=-.298, *p*=.025, n = 56), norcocaine (r=-.302, *p*=.023, n = 56), cocaine (r=-.278, *p*=.037, n = 56) metabolites, and Cocaine_total_ – the sum of the cocaine metabolite, benzoylecgonine metabolite and norcocaine metabolite – (r=-.291, *p*=.029, n = 56) on *risk* sensitivity, indicating that a severe cocaine consumption pattern goes along with a lower risk sensitivity. Nevertheless, these effects did not remain significant after including IQ, executive functioning, and ADHD symptoms in the model, in accordance with the hierarchical linear models (Table 3). Regarding *gain* and *loss* sensitivity, we found no effect for the self-report estimated cumulative lifetime dose nor cocaine metabolites with and without including years of school education in the models (see Table 3).

## 4. Discussion

Our study extends current knowledge on decision-making deficits in CU by analysing risky decisions in the CCT with higher resolution. We investigated whether chronic CU differed from stimulant-naïve controls in the use of *gain, loss, and risk* information during decision-making under risk. CU were more risk-seeking than controls in less favourable decision scenarios, where returns were low and risk was high (i.e., lower expected value). By looking at the use of information over all decision scenarios, the data confirmed our hypothesis that chronic CU are not as sensitive to *gains* as stimulant-naïve controls. Indeed, CU were less sensitive to the expected value, suggesting that they are not able to fully integrate all the available information. We also found a marginally significant group effect for *loss* sensitivity; however, no group effect was found for *risk* sensitivity. Furthermore, the main group difference in *gain* sensitivity was not explained by additional predictors (i.e., IQ, executive functioning, working memory, visual processing efficiency, impulsivity traits nor ADHD symptoms), although years of school education also had an effect. By contrast, *loss* sensitivity was related to years of education, but not group and, for *risk* sensitivity, we found an effect for IQ, executive functioning and ADHD symptoms, but not with group. Finally, the correlation analyses between risk-attitude and the sensitivity to *gain, loss, and risk* showed that chronic CU are, unlike stimulant-naïve controls, less able to consider all available information on returns (i.e., gain and loss) and risk. From a clinical perspective, the lack of gain sensitivity is not surprising, since one of the core criteria of all substance-related disorders is the withdrawal from social, occupational, and recreational activities with high value in order to use the substance [52]. This pattern of behaviour was also proposed to reflect a shift in the subjective value of ordinary life events to substance-related rewards [23].

Concerning *loss* and *risk* information, we found a significant effect of years of school education on *loss* sensitivity and a significant effect of verbal IQ, executive functioning, and ADHD symptoms on *risk* sensitivity, but no effect of group status per se. Thus, in contrast to *gain* sensitivity, which was clearly altered in the CU group, *loss* and *risk* sensitivity are better explained by demographic and intellectual differences as well as psychiatric comorbidities than by chronic cocaine use. These results demonstrate that the interpretation of deficits in decision-making findings needs to take into account the particular demographic and clinical background [53] typically associated with cocaine-related disorder. Given that within the CU group 90% met the criteria for current or past cocaine dependency or abuse according to DSM-IV-R, we expected to find higher self-reported impulsivity and ADHD symptoms and worse general cognitive performance in the CU than the control group. Hence, in our study, the stimulant-naïve control group served as a normative sample and was matched for sex, age, smoking status, and current pattern of alcohol consumption, while differences in IQ, years of school education, self-reported and behavioural impulsivity, ADHD symptoms, and cognitive performance were all considered when performing the hierarchical linear models. Although our findings suggest that severe cocaine use is not directly linked to a decrease in sensitivity to *loss* and *risk* information, it does show that typical cocaine users, nevertheless, may display impairments in the processing of loss and risk information.

Our data also suggest that CU may not as fully integrate all the available information as controls when making risky decisions, as shown by the interaction effect of group and expected value on risky behaviour. Such finding was supported by the correlations between information sensitivity and risk-attitude, which showed that within the control group, sensitivity to *gain* and *loss* information negatively correlates with increased risky behaviour and sensitivity to *risk* information positively correlates with increased risky behaviour in the most unfavourable decision scenario; while in the most favourable decision scenario sensitivity to *gain* and *loss* information positively correlates with increased risky behaviour and sensitivity to *risk* information negatively correlates with increased risky behaviour. However, within the CU group, we found that in the most unfavourable decision scenario, only sensitivity to *gain* and *loss* information negatively correlates with increased risky behaviour, but no correlation with sensitivity to *risk* information was found; while in the most favourable decision scenario, only sensitivity to *risk* information negatively correlates with increased risky behaviour, but no correlation for sensitivity to *gain* and *loss* information was found. Such impairments in integrating all the available information could be related to vmPFC dysfunction, as this brain region has been associated with the integration of subcortical signals into a single representation of net value, which is accumulated over time until the individual decides to accept or reject an option [54]. Indeed, in the Iowa Gambling Task, CU showed impaired performance that resembled the maladaptive behaviour of patients with vmPFC lesions [55, 56]. More specifically, CU also showed reduced vmPFC activation to social and object reward [27], in line with a *gain* processing function of this region and the reduced *gain* sensitivity of CU found here.

Some limitations should be considered when interpreting our findings. First, the cross-sectional design of this study does not allow us to clearly determine the causal relationship between cocaine use and alterations in *gain* sensitivity, especially because we found no correlation with subjective and objective cocaine use severity markers. Thus, it’s also plausible that a lower sensitivity to *gain* predicts the onset of substance use. Nevertheless, it seems to be more likely that variations in *loss* and *risk* sensitivities precede chronic cocaine use, as indicated by the significant effects of demographic, clinical, and cognitive variables. Future studies might consider investigating whether changes in cocaine consumption can affect sensitivity for *gain*, *loss*, and *risk* information during decision-making. Remarkably, this finding complements one of our previous studies that investigated decision-making under risk without feedback using a different definition of risk and found that risk proneness was associated with higher cocaine concentrations in the hair [57]. The different definition of risk may also explain why, in contrast to Wittwer, et al. [57], we found an effect of IQ, executive functioning, and ADHD symptoms on *risk* sensitivity, but no effect for sex and age.

Taken together, our findings open avenues for future applied research that aims to improve the efficiency and the efficacy of preventive and therapeutic strategies for chronic substance users. For instance, decreased sensitivity to *gain* might partially explain the lack of adherence to long term treatments and detoxification programs, since chronic CU are insensitive to the possible long-term advantages of maintaining abstinence. In addition, our findings support the necessity of considering demographic, clinical, and cognitive variables when providing therapeutic strategies, something that is well-known, but frequently not applied.

## Acknowledgements

We are grateful to Monika Visentini, Monika Näf, Chantal Kunz, Selina Maisch, Marlon Nüscheler, Anna Burkert, Meret Speich, Maxine de Ven, Jocelyn Waser, Zoe Dolder, Zoe Hillmann, Jessica Grub, and Priska Cavegn for all their effort in data collection.

## Ethical standards

The authors assert that all procedures contributing to this work comply with the ethical standards of the relevant national and institutional committees on human experimentation and with the Helsinki Declaration of 1975, as revised in 2008.

