## Supplement for "Sensitivity to gains during risky decision-making differentiates chronic cocaine users from stimulant-naïve controls"

### Supplementary Material

### Materials and Methods

##### Exclusion criteria

Exclusion criteria comprised a family history of genetically mediated psychiatric disorders ($h^2$>0.5, e.g., autism, schizophrenia, and bipolar disorder); any severe neurological disorder or brain injury; a current diagnosis of infectious diseases or severe somatic disorder; a history of autoimmune, endocrine, and rheumatoid arthritis; intake of medication with potential action at the central nervous system during the last three days; participation in a large previous study from our lab, the *Zurich Cocaine Cognition Study* ($ZuCo^2St$) (Vonmoos. *et al.*, 2014); and for women being pregnant or breastfeeding. Controls were excluded if they had DSM-IV-R Axis I adult psychiatric disorders, or recurrent illegal substance use (>15 occasions lifetime, with the exception of cannabis due to reasons of participant matching). We excluded CU with regular use of illegal substances other than cocaine such as heroin or other opioids (with exception of cannabis use), a polysubstance use pattern according to DSM-IV-R, or a DSM-IV axis I adult psychiatric disorder diagnosis (e.g., schizophrenia, bipolar disorder, current major depressive episode, eating disorders, current anxiety disorder) except for cocaine, cannabis, and alcohol abuse/dependence, previous depressive episodes, and ADHD.

##### Procedure

Data were collected at the Psychiatric Hospital of the University of Zurich. CU were recruited from treatment centres for addiction in and around the Canton of Zurich, Switzerland. The testing sessions started either at 09:00am or noon and lasted for around five hours. Participants were asked to abstain from illegal substances for a minimum of 72h and from alcohol for at least

24h before the session. Each participant was tested individually and anonymously. All clinical and cognitive assessments were performed by trained psychologists or students in psychology who were supervised by a clinical psychologist.

#### Hair toxicological analysis

A total of 98 compounds and metabolites were accessed, as shown in Table S2. For a complete description of our routine protocol and methodological adaptations, please see Vonmoos et al. (2013), Scholz et al. (2019), and Scholz et al. (submitted) respectively.

**Table S2.**
Complete list of compounds (metabolites) accessed.

| Class | Drug (metabolite) |
|---|---|
| Analgesic/Migraine | Diclofenac<br>Metamizole<br>Naratriptan<br>Paracetamol<br>Sumatriptan |
| Anticonvulsants | Flunarizin<br>Gabapentin<br>Lamotrigin<br>Pregabalin |
| Antidepressants | Agomelatine<br>Amitriptyline<br>Bupropione<br>Citalopram<br>Clomipramine<br>Doxepine<br>Duloxetine<br>Fluoxetin<br>Fluvoxamine<br>Imipramine<br>Mirtazapine<br>Nortriptyline<br>Opipramol<br>Paroxetine<br>Sertraline<br>Trazodone<br>Trimipramine<br>Venlafaxine (O-Desmethylvenlafaxine) |
| Antihistamines | Diphenhydramine<br>Doxylamine |
| Antipsychotics | Amisulpride<br>Asenapine<br>Chlorprothixene<br>Clozapine<br>Haloperidol<br>Levomepromazine<br>Olanzapine<br>Paliperidone<br>Pipamperone<br>Promazine<br>Quetiapine (Norquetiapine, OH-Quetiapine)<br>Risperidone (OH-Risperidone) |
| Benzodiazepines | Alprazolam<br>Bromazepam<br>Demoxepam<br>Diazepam<br>Clobazam (N-Desmethylclobazam) |

| | |
|---|---|
| | Clonazepam (7-Aminoclonazepam) |
| | Flunitrazepam (7-Aminoflunitrazepam) |
| | Flurazepam (N-Desalkylflurazepam) |
| | Lorazepam |
| | Midazolam (α –Hydroxymidazolam) |
| | Nitrazepam (7-Aminonitrazepam) |
| | Nordazepam |
| | Oxazepam |
| | Phenazepam |
| | Prazepam |
| | Temazepam |
| | Tetrazepam |
| | Triazolam |
| Cannabinoids | Cannabidiol |
| | Cannabinol |
| | Tetrahydrocannabinol |
| Entactogens | 2,5-Dimethoxy-4-bromophenethylamine, 2C-B |
| | 4-Fluoroamphetamine, 4-FA |
| | 3,4-Methylenedioxy-N-ethylamphetamine (MDEA) |
| | 3,4-Methylenedioxymethamphetamine, MDMA (MDA) |
| Opioids | Acetylmorphine |
| | Acetylcodeine |
| | Buprenorphine (Norbuprenorphine) |
| | Codeine |
| | Dihydrocodeine |
| | Fentanyl (Norfentanyl) |
| | Hydromorphone |
| | Methadone (1,5-Dimethyl-3,3-diphenylpyrrolidine (EDDP)) |
| | Morphine |
| | Oxycodone |
| | Oxymorphone |
| | Pethidine |
| | Tapentadol |
| | Tilidin |
| | Tramadol (N-Desmethyltramadol) |
| Other substances | Levamisole |
| | Dextromethorphan |
| | Clomethiazole |
| | Tizanidine |
| | Ketamine (Norketamine) |
| Stimulants | Amphetamine |
| | Atomoxetine |
| | Cocaine (Benzoylecgonine, Cocaethylene, Norcocaine) |
| | Methamphetamine |
| | Methylphenidate |
| | Modafinil |
| Z-Hypnotics | Zaleplon |
| | Zolpidem |
| | Zopiclone |

#### General cognitive assessment

In addition to the estimate of premorbid verbal intelligence measured by the German vocabulary test *Mehrfachwahl-Wortschatz-Intelligenztest* (Lehrl, 1999), general cognitive performance was assessed with a selection of three tasks from the *Cambridge Neuropsychological Test Automated Battery* (CANTAB, http://www.cantab.com): the *Spatial Working Memory* task (SWM), assessing working memory and executive functioning; the *Match to Sample Visual Search* task (MTS), a visual matching test involving a speed-accuracy trade-off; and the *Rapid Visual Information Processing* task (RVP), assessing sustained attention capacity.

The *Spatial Working Memory* task (SWM) was used to assess spatial working memory and executive functioning. In this task a number of coloured boxes are shown on the screen. Participants should find a yellow 'token' in one of the boxes and use them to fill up an empty column on the right-hand side of the screen. Depending on the difficulty level used for this test, the number of boxes can be gradually increased until a maximum of 12 boxes. We measured between-search errors (occasions the participant returned to search a box in which a token had already been found during a previous search sequence), within-search errors (occasions the participant revisited a box already found to be empty during the same search sequence), total errors, and a strategy score (number of times the participant began a new search with a different box to the last search; therefore, a high score indicated an inefficient strategy).

The *Match to Sample Visual Search* task (MTS) is a visual matching test involving a trade-off of speed and accuracy. In this task the sample stimulus is an abstract pattern displayed within a red square in the middle of the screen. After a brief delay, a varying number of similar patterns (1, 2, 4 or 8) is shown in a circle of boxes around the edge of the screen. Only one of these patterns matches the pattern in the centre of the screen. The subject must select the matching stimulus by touching it. The task provides three major measures: the percentage of correct responses, the mean latency of correct responses (the time taken to respond to trials correctly), and the mean change in movement time between trials with 2 and 8 choices (mean movement time in trials with 8 choices minus trials with 2 choices).

The *Rapid Visual Information Processing* task (RVP) was designed to assess sustained attention capacity. In this task, a white box appears in the centre of the screen, inside which single digits appear in a pseudo-random order, at the rate of 100 digits per minute. Subjects are requested to detect target sequences of digits (for example, 2-4-6, 3-5-7, 4-6-8) and to register responses using the press pad. The task provides the following measures: total false alarms (occasions the participant responded inappropriately), response A' (a signal detection measure of the sensitivity to the target) response bias B' (a signal detection measure of the strength of trace required to elicit a response), and mean latency for correct responses (mean time taken to respond correctly).

Moreover, based on our previous work, we calculated the *Impulsivity index* and the *Efficiency index* for the MTS and RVP (Quednow *et al.*, 2007, Vonmoos *et al.*, 2013). The

Impulsivity index is a score quantifying the behavioural dimension of “fast and inaccurate” vs. “slow and accurate”. For the RVP, this score was obtained by subtracting the z-standardized score of the mean latency over all responses from the z-standardized score of the total of false alarms, while for the MTS, this index was obtained by subtracting the z-standardized score of the overall reaction time from the z-standardized score of the total of errors committed. The Efficiency index is a score quantifying the “fast and accurate” vs. “slow and inaccurate” dimension, and was obtained from the RVP by summing the z-standardized score of the total of false alarms committed and the z-standardized score of the mean latency over all responses and multiplying the sum with –1. Similarly, for the MTS, this score was obtained by summing the standardized score of the total of errors committed and the standardized score of the overall reaction time and multiplying the sum with –1. The false alarms and latency values from the RVP and the errors committed and the reaction time from the MTS of all participants were standardized to the means and standard deviations of the control group. The Meta-Efficiency Index was calculated by taking the average of both Efficiency indices. Previous work from our group has shown that, in a visual search task, a similar impulsivity index can distinguish MDMA users from controls (Quednow *et al.*, 2007), although in the RVP task we found no impulsivity index differences between cocaine users and stimulant-naïve controls (Vonmoos *et al.*, 2013). Finally, a Meta-Efficiency Index was calculated by taking the average across the Efficiency indices of both the MTS and the RVP.

#### The Columbia Card Task

In the no-feedback version of the CCT participants were explicitly instructed to perform each round independently from the previous one, because at the end of the task three rounds would be randomly selected to calculate the payoff. The eight decision scenarios were repeated three times and randomly presented within three blocks, for a total of 24 rounds. In this version of the CCT, participants did not receive any feedback about their performance until the end of the 24 rounds, when three rounds were randomly selected by the computer and the sum of the points made in these rounds was converted to actual money for the participant (1 point = 0.10 CHF).

##### *Risk-attitude*

Given that both the gain and the likelihood of experiencing a loss increased with each card turned over by the computer, choosing more cards to be turned over is associated with greater outcome variability and, therefore, was a riskier strategy than turning over fewer cards. Thus, in the risk-attitude analysis we determined risk-seeking propensity independently from the influence of *gain*, *loss,* and *risk*.

### Results

**Table S1.**
Spearman's rank correlations between all predictors included in the hierarchical multiple linear regression models.

| | Years of Education | IQ | SWM Strategy | SWM Total Errors | Efficiency Index [a] | BIS | ADHD |
|---|---|---|---|---|---|---|---|
| Years of education | 1 | - | - | - | - | - | - |
| IQ | .261 ** | 1 | - | - | - | - | - |
| SWM Strategy | **-.307** ** | -.133 | 1 | - | - | - | - |
| SWM Total Errors | -.233 * | -.166 | **.774** *** | 1 | - | - | - |
| Efficiency Index [a] | .173 | .125 | **-.389** *** | **-.372** *** | 1 | - | - |
| Impulsivity Index [a] | -.120 | .029 | .088 | .044 | -.012 | - | - |
| BIS | -.199 | -.042 | .173 | .117 | -.095 | 1 | - |
| ADHD | -.162 | -.037 | .176 | .100 | -.163 | **.497** *** | 1 |

Note. Correlations were performed within the total sample (n = 96). YE, Years of Education; IQ, Intelligence Quotient assessed with the Mehrfachwahl-Wortschatz-Intelligenztest (MWT-B); EF, Executive Functioning assessed with the Strategy score from the Spatial Working Memory task (SWM); WM, Working Memory assessed with the Total Errors from the Spatial Working Memory task; BIS, total sum score from Barratt Impulsiveness Scale; ADHD, Attention deficit hyperactivity disorder total sum score from ADHD self-rating scale. $p$-value $\leq .05$ *; $p$-value $< .01$ **; $p$-value $< .001$ ***. (a) Efficiency Index is the average of the efficiency indices from the Rapid Visual Information Processing task and the Match to Sample Visual Search task.
